# Ecology drives the degree of convergence in the gene expression of extremophile fishes

**DOI:** 10.1101/2021.12.13.472416

**Authors:** Michael Tobler, Ryan Greenway, Joanna L. Kelley

## Abstract

Convergent evolution, where independent lineages evolve similar traits when adapting to similar habitats, is a common phenomenon and testament to the repeatability of evolutionary processes. Still, non-convergence is also common, and a major question is whether apparently idiosyncratic, lineage-specific evolutionary changes are reflective of chance events inherent to evolutionary processes, or whether they are also influenced by deterministic genetic or ecological factors. To address this question, we quantified the degree of convergence in genome-wide patterns of gene expression across lineages of livebearing fishes (family Poeciliidae) that span 40 million years of evolution and have colonized extreme environments in the form of toxic, hydrogen-sulfide-rich springs. We specifically asked whether the degree of convergence across lineage pairs was related to their phylogenetic relatedness or the ecological similarity of the habitats they inhabit. Using phylogenetic comparative analyses, we showed that the degree of convergence was highly variable across lineage pairs residing in sulfide springs. While closely related lineages did not exhibit higher degrees of convergence than distantly related ones, we uncovered a strong relationship between degree of convergence and ecological similarity. Our results indicate that variation in the degree of convergence is not merely noise associated with evolutionary contingency. Rather, cryptic environmental variation that is frequently ignored when we employ reductionist approaches can significantly contribute to adaptive evolution. This study highlights the importance of multivariate approaches that capture the complexities of both selective regimes and organismal design when assessing the roles of determinism and contingency in evolution.

**Significance Statement:** When different species adapt to similar environmental conditions, we frequently observe a mix between shared (convergent) and lineage-specific (nonconvergent) evolutionary changes. Shared changes provide evidence for the repeatability and predictability of evolution. However, it remains unclear whether lineage-specific changes are caused by random forces that limit the predictability of evolution, or whether they reflect deterministic processes shaped by unidentified genetic and environmental factors. By analyzing patterns of gene expression across fishes in extreme environments, we show that the degree of convergence between lineages is related to ecology, indicating that lineage-specific evolutionary changes are not just noise caused by random processes. Thus, acknowledging the complexity of nature in empirical research is critical if we want to predict evolution.

## Introduction

A core question in contemporary biology is whether evolution repeats itself, and—if so—whether we can predict the circumstances under which repeatable evolutionary change happens (1). The question of evolutionary repeatability is epitomized by the contrasting perspectives of two giants in the field. On one side, Stephen J. Gould was adamant about the role of contingency in evolution, famously concluding that the “tape of life” would never repeat itself due to chance events inherent to the evolutionary process (2). On the other side is Simon Conway Morris, a strong proponent of evolutionary determinism, who likened evolutionary change to a drop of water that follows the pull of gravity, flowing down a landscape according to a predictable path of least resistance, and inevitably ending up in the ocean. Upon investigating much of the same evidence as Gould, Conway Morris argued that evolution does repeat itself, because there are a finite number of engineering solutions to the problems posed by a finite set of niches available to life on Earth (3).

Past research has unearthed much evidence for both contingency and determinism in evolution. The importance of random mutation and genetic drift in evolutionary change are just as undeniable (4) as the prevalence of convergent evolution (5–7), where independent lineages evolve similar solutions to shared ecological problems. Hence, the question is not so much whether chance events happen, or whether evolution can repeat itself, but instead whether we can predict the circumstances under which determinism prevails over contingency. At the root of this problem are confirmation biases inherent to the detection of deterministic and contingent evolutionary outcomes. It is relatively easy to document the presence of convergence (8–10). However, a lack of convergence is not necessarily evidence for a role of contingency in evolution; failure to readily understand the origins of lineage-specific idiosyncrasies does not exclude deterministic explanations in favor of invoking contingency. In fact, unique, lineage-specific evolutionary outcomes in response to apparently similar selective regimes can arise through a multitude of genetic, functional, and ecological mechanisms (11). For example, differences in genetic architecture can bias responses to selection (12), functional redundancy can cause equivalent fitness by alternative means (13), and cryptic ecological differences among lineages can select for different phenotypic optima (14). Still, attempts to explain the degree of convergent and lineage-specific (*i.e*., nonconvergent) evolutionary patterns remain scarce.

We leveraged a unique natural experiment to test whether we can explain variation in the degree of convergent evolution in response to a strong and clearly defined source of selection. Throughout the Neotropics, fish of the family Poeciliidae have colonized springs rich in hydrogen sulfide (H_2_S) (15), a potent respiratory toxicant lethal to most metazoans in micromolar concentrations (16). Sulfide spring lineages span over 40 million years of evolutionary divergence, facilitating comparisons between populations of the same species and between species within and among genera (Figure 1A). Previous research has documented that sulfide spring lineages show clear signatures of convergent evolution of mitochondrial genomes, the regulation of genes associated with H_2_S toxicity and detoxification, and aspects of physiological function, morphology, reproductive life history, and behavior (17–21). Despite the evidence for convergence across levels of organismal organization and across a wide gradient of phylogenetic relationships, past studies have also found evidence of lineage-specific evolutionary change, giving rise to substantial variation in the degree of convergent evolution between specific pairs of sulfide spring lineages (22).

**Figure 1.**
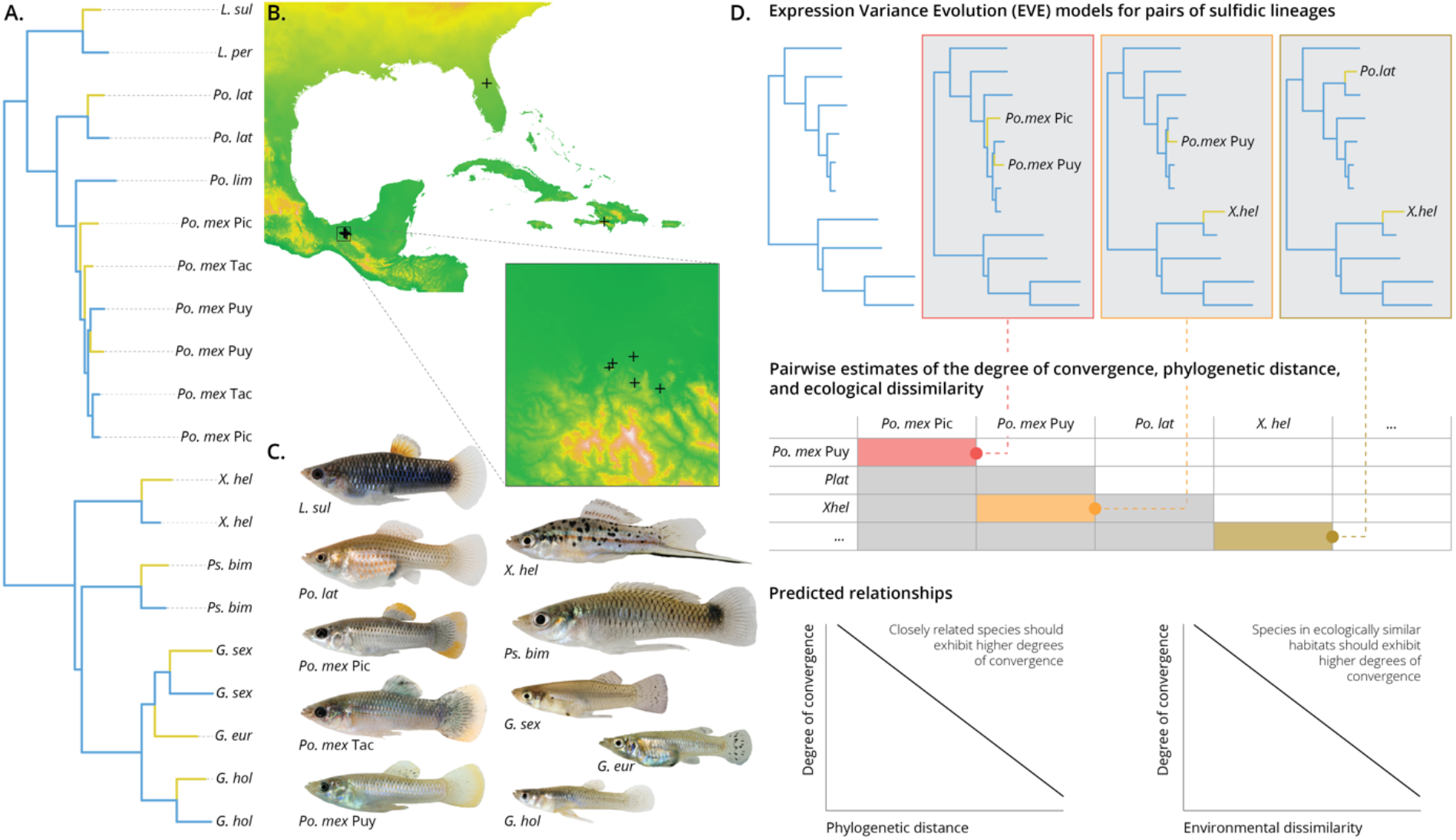
A. Phylogeny of lineages included in this study. Yellow edges indicate lineages that have colonized sulfide springs. B. Map of the locations of different sulfide springs. Samples were collected from Florida (one spring), the Dominican Republic (one spring) and Mexico (five springs; region enlarged in inset). C. Representative specimens of the sulfide spring lineages investigated here: *Limia sulphurophila* (*L. sul*); *Poecilia latipinna* (*Po. lat*); *Po. mexicana* (*Po. mex*) from the Ríos Pichucalco (Pic), Tacotalpa (Tac), and Puyacatengo (Puy); *Xiphophorus hellerii* (*X. hel*); *Pseudoxiphophorus bimaculatus* (*Ps. bim*); *Gambusia sexradiata* (*G. sex*); *G. eurystoma* (*G. eur*); and *G. holbrooki* (*G. hol*). D. Outline of the analytical framework used in this study. A phylogeny of all nonsulfidic lineages served as a backbone for all analyses, allowing to account for gene expression variation that arose as a consequence of the diversification of lineages. To identify convergent shifts in gene expression upon the colonization of sulfide springs, sulfidic lineages were added to the backbone phylogeny in a pairwise fashion (as exemplified by the red, orange, and brown phylogenies) to run expression variance and evolution (EVE) models. The output of EVE models provided quantitative measures of the degree of convergence between pairs of sulfide spring lineages, which were complemented by estimates of phylogenetic divergence and ecological dissimilarity. The degree of convergence between pairs of sulfide spring populations was predicted to be negatively correlated both with their phylogenetic distance and the ecological dissimilarity of their habitats.

Here, we explicitly tested whether we could explain variation in the degree of convergent evolution among 10 sulfide spring populations of the family Poeciliidae, including lineages in the genera *Limia*, *Poecilia*, *Xiphcphorus*, *Pseudoxiphophorus*, and *Gambusia* from three different biogeographic regions (Figure 1A–C). Specifically, we used phylogenetic comparative methods to analyze genome-wide patterns of gene expression and asked whether convergence in gene expression between pairs of sulfide spring populations was dependent on their phylogenetic relationship or the ecological similarity of their habitats. We predicted that more closely related species would exhibit higher degrees of convergence (23), because shared genomic architectures would determine similar responses to selection (Figure 1D). In addition, we predicted that species exposed to more similar ecological conditions would exhibit higher degrees of convergence (14), because environmental variables that covary with the presence of H_2_S would determine the trajectories of evolutionary change (Figure 1D). If neither of these predictions held true, then among-lineage variation may indeed be caused by contingencies.

## Results

We analyzed 118 gill transcriptomes from ten lineages of sulfide spring poeciliids and ten closely related lineages from adjacent nonsulfidic habitats (17). To quantify variation in the degree of convergence in gene expression between pairs of sulfide spring populations, we selected the 5,000 top expressed genes across all lineages as dependent variables in expression variance and evolution (EVE) models (24, 25), which allow for the identification of convergent expression shifts while accounting for the evolutionary relationship among lineages [see Figure 1A; (26)]. EVE models implement extended Ornstein-Uhlenbeck processes accounting for within-lineage variation to test for shifts in gene expression between a set of background lineages (populations from nonsulfidic habitats) and a set of foreground lineages (populations from sulfidic habitats). Models were run separately for all pairwise combinations of sulfidic lineages (a total of 45 pairwise comparisons), such that each model included ten nonsulfidic lineages as the background and two sulfidic lineages as the foreground (Figure 1D). Hence, genes with convergent expression shifts for each sulfidic population pair were identified under explicit consideration of macroevolutionary patterns of gene expression, and inference of convergence for a particular population pair was not biased by the presence of other sulfidic lineages in the analysis.

The number of convergent gene expression shifts (FDR < 0.05) identified through this approach ranged from 1 (between *L. sulphurophila* and *P. mexicana* Puy) to 308 (between *G. eurystoma* and *X. hellerii*, Figure 2A, Table S1). Consistent with theoretical predictions (16) and previous analyses (17), the genes with evidence for convergent expression shifts in the majority (≥ 23) of comparisons were associated with H_2_S detoxification, the processing and transport of sulfur compounds, oxidative stress responses, and redox homeostasis (Figure S1). In contrast, genes with evidence for convergent expression shifts in a minority of comparisons were primarily associated with the organization of mitochondria and the assembly of the mitochondrial respiratory chain [particularly complex IV, which is the primary toxicity target of H_2_S (27)], aerobic and anaerobic metabolism, as well as ion and mitochondrial transmembrane transport (Table S2).

**Figure 2.**
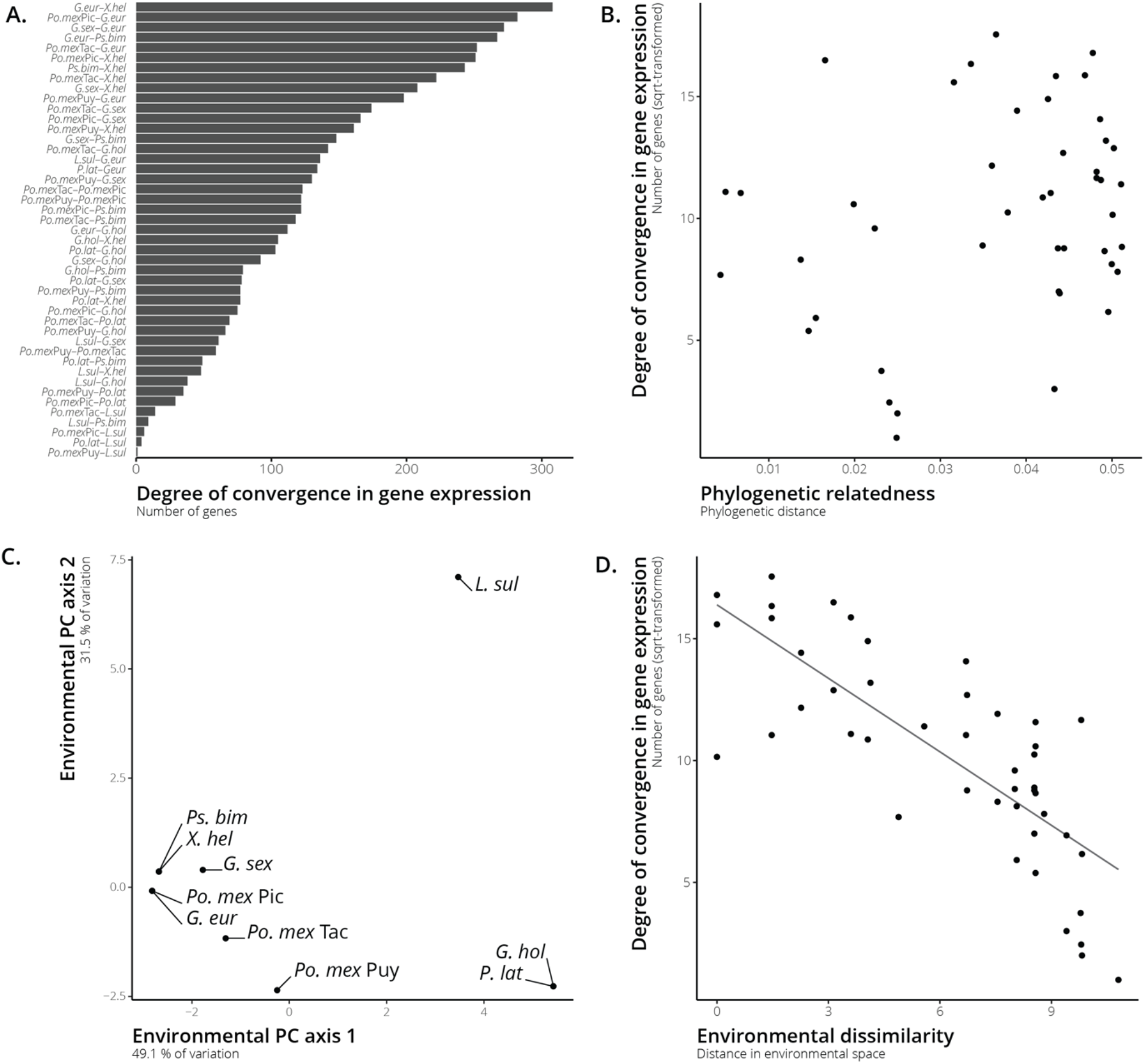
A. Variation in the degree of convergence in gene expression among all pairs of lineages from sulfidic environments spanned over two orders of magnitude. B. Variation in the degree of convergence was not correlated with phylogenetic relatedness (*P* = 0.977). C. Visualization of environmental variation among different sulfide springs based on a principal component (PC) analysis. PC axis 1 was primarily correlated with pH, water temperature, and H_2_S concentrations, as well as bioclimatic variables 1, 4, 6, 7, 9, 11, 12, 13, 14, 16, 17, 18, and 19 (see Table S1 for details). Note that some springs harbor more than one lineage of H_2_S-adapted fish. D. Variation in the degree of convergence between different sulfide spring lineages was strongly correlated with the environmental dissimilarity between their habitats (*r* = −0.781, *P* < 0.0001).

To test whether more closely related species exhibited higher degrees of convergence, we calculated pairwise phylogenetic distances between all lineages from sulfidic habitats and used a partial Mantel test to assess the correlation between the degree of convergence and the degree of evolutionary relatedness. Contrary to our prediction, the degree of convergence in gene expression exhibited no relationship with phylogenetic distance (Figure 2B; partial Mantel: *r* = 0.289; *P* = 0.977). Hence, there was no evidence that closely related lineages exhibit higher degrees of convergent evolution than distantly related ones.

To test whether population pairs in more ecologically similar habitats exhibit higher degrees of convergence, we measured aspects of water quality *in situ* (temperature, pH, specific conductivity, dissolved oxygen, and H_2_S concentration) and downloaded 19 geospatial variables that describe climatic variation for each sulfide spring (28). Principal component (PC) analysis indicated that different springs primarily segregated along the first PC axis (Figure 2C), which was correlated with temperature, pH, and H_2_S concentration, as well as bioclimatic variables 1, 4, 6, 7, 9, 11, 12, 13, 14, 16, 17, 18, and 19 (correlations: |0.20| < *r* < |0.28|; Table S1). In general, sulfide springs in Mexico (negative scores along PC axis 1) exhibited higher temperatures and H_2_S concentrations, lower pH, less seasonality, and more precipitation compared to the springs in Florida and the Dominican Republic (positive scores along PC axis 1). Variation along the second PC axis was primarily driven by a single outlier, the sulfide spring from the Dominican Republic. To quantify ecological dissimilarity between different sulfide spring lineages, we calculated pairwise Euclidean distances in multivariate environmental space (24 dimensions based on *z*-transformed variables), and again used a partial Mantel test to assess correlation between the degree of convergence and ecological dissimilarity. As predicted, we detected a strong negative correlation between the degree of convergence and ecological dissimilarity across different pairs of sulfide spring populations (Figure 2B; partial Mantel: *r* = −0.781; *P* < 0.001). Hence, the more similar ecological conditions were in the habitats of the focal lineages, the higher the number of genes with evidence for convergent expression shifts.

## Discussion

Our phylogenetic comparative analyses of gene expression among ten lineages of sulfide spring fishes revealed starkly different patterns of convergence. The number of genes with evidence for convergent expression shifts varied over two orders of magnitude across different lineage pairs. Genes with evidence for convergent expression shifts in the majority of lineage pairs were particularly associated with H_2_S detoxification, including 2-mercaptopyruvate sulfurtransferase (MPST), persulfide dioxygenase (ETHE1), and sulfide:quinone oxidoreductase (SQRDL), all of which play critical roles in the enzymatic oxidation of H_2_S to nontoxic sulfur species (29). Genes with evidence for convergent expression shifts in a minority of lineage pairs were particularly associated with the function of the mitochondrial respiratory chain and the modulation of aerobic and anaerobic metabolism. This finding is not surprising considering that H_2_S primarily unfolds its toxicity by interrupting aerobic ATP production in mitochondria, as it binds reversibly to complex IV of the respiratory chain (27). Overall, our results are consistent with previous studies on a much smaller phylogenetic scale, which showed that different sulfide spring lineages of *Po. mexicana* consistently modulate genes associated with H_2_S detoxification (19), but only a subset of lineages also exhibits modifications to the respiratory chain that reduce the impact of H_2_S toxicity (30).

Perhaps most importantly, we found clear evidence that the degree of convergence among sulfide spring lineages is related to the ecological similarity of their habitats. Lineage-specific idiosyncrasies in evolutionary change are thus unlikely to be mere consequences of contingencies. Rather, cryptic environmental variation, which is ignored when habitats are just classified based on the presence and absence of H_2_S (11), must influence the evolutionary trajectories of lineages in a consistent way. The importance of cryptic environmental variation in shaping the degree of evolutionary convergence was also stressed by a study of replicated pairs of threespine stickleback (*Gasterosteus aculeatus*) that inhabit lake and stream habitats (14). Like in our study, the degree of convergence among population pairs was highly variable and related to differences in habitat structure, trophic resource use, and parasite communities (14). Our study highlights that the importance of cryptic environmental variation does not only shape the degree of convergent evolution among closely related populations of the same species, where selection likely acts on shared pools of standing genetic variation, but it also does so at much broader phylogenetic scales.

If Gould was able to weigh in at this stage, he might caution us not to fall victim to the adaptationist fallacy (31); after all, correlation does not imply causation. Clearly, future research needs to address how variation in complex organismal traits impacts the fitness of individuals in the context of the multifarious selective regimes found in sulfide springs. Unfortunately, this is not trivial empirically. While strong patterns of convergence provide tangible candidate traits to establish the relationships between form, function, and fitness, the field of potential candidates is much wider when we consider traits that are modified in just one or a subset of lineages. This is a general problem in evolutionary analyses, and a major question moving forward is how we can be more explicit about the complexity of environments and organisms when studying evolutionary patterns and processes. Gaining a fuller picture about the function of organisms in their natural habitat will require us to move beyond reductionist approaches and consider that lineage-specific evolutionary patterns are not just noise but a consequence of deterministic forces and play a role in adaptation.

A major gap that remains unaddressed by our study is to what degree patterns of convergence in gene expression documented here are the product of evolutionary change in gene regulatory mechanisms as opposed to shared plastic responses to the environmental conditions that sulfide spring fishes encounter in their habitats. Studies in *Po. mexicana* have indicated that ancestral plasticity plays a minor role in shaping gene expression differences between replicated sulfidic and nonsulfidic populations (32). Instead, population differences in gene expression have a strong genetic basis and are caused by changes in both the constitutive expression and the H_2_S-inducibility of genes associated with adaptation to sulfidic conditions (32). The evidence for evolved changes in gene regulation at a much smaller phylogenetic scale—together with the high heritability of genome-wide mRNA expression patterns documented in other systems (33, 34)—therefore suggests that the patterns of convergence across sulfide spring lineages are not merely a consequence of shared plastic responses but are at least in part shaped by adaptive evolution. Future studies still need to undertake the daunting task of assessing the heritability in gene expression variation beyond *Po. mexicana*, although the strong effect of cryptic ecological variation on the degree of gene expression similarity across lineages is remarkable even if some of the shared responses are a consequence of plasticity.

Our study also revealed that phylogenetic relatedness did not predict the degree of convergence among different sulfide spring lineages. Closely related species are typically assumed to have a similar genetic architecture and may even share patterns of genetic variation, leading to similar evolutionary responses when exposed to the same sources of selection (23). One interpretation of our finding is that constraints associated with genetic variation and genomic architecture do not play a significant role in shaping responses to selection in sulfide springs. However, it is important to note that H_2_S interacts with ancient physiological pathways that are shared across metazoans. H_2_S is toxic because it interferes with oxidative phosphorylation, a pathway that has been highly conserved throughout the evolution of life (35, 36). Similarly, H_2_S is detoxified through the mitochondrial sulfide:quinone oxidoreductase pathway, which has prokaryotic origins and is highly conserved in metazoans (37, 38), although the detoxification capacity in most species is limited to endogenously produced H_2_S and rapidly overwhelmed when exposed to environmental H_2_S (39). Consequently, adaptation to sulfide springs likely did not require the evolution of novel genes or gene functions, but merely the modification of ancient pathways that were already shared among all lineages. In addition, the clear-cut biochemical and physiological consequences of H_2_S may have constrained evolutionary responses to a few pathways, such that among-lineage variation in genetic variation and genomic architecture did not bias responses to selection substantially. Insights from other study systems where responses to selection are less constrained will be critical to test the general role of phylogenetic distance in shaping the degree of convergent evolution.

Finally, there appears to be a chasm between patterns of convergent evolution at a genomic level and emerging phenotypic traits. Convergence in phenotypes is common, but it is comparatively rare at the genomic level (10), unless constraints are substantial (40), selection acts on standing genetic variation (41), or adaptive alleles are introgressed across divergent lineages (42). This chasm is also evident in the sulfide spring fishes studied here. Specifically, *Po. mexicana* (Pic), *Ps. bimaculatus*, and *X. hellerii*—which all inhabit the same sulfide spring—exhibit high levels of convergence in gene expression (Figure 2A) and other traits (43). Yet population genomic analyses revealed only scant evidence for convergence at a genomic level, and none of the outlier loci were associated with genomic regions containing genes associated with H_2_S toxicity and detoxification (43). These contrasting results should not be surprising, and to some degree they reconcile the conflicting views Gould and Conway Morris had on the role of contingency and determinism in evolution. We have known since Kimura’s (44) introduction of the neutral theory that drift profoundly shapes genetic variation within and between species. More so, redundancy in the genetic basis of quantitative traits involved in adaptation can cause idiosyncratic responses to selection that are in part influenced by chance (45, 46). Hence, genomic evolution largely reflects Gould’s notion of the importance of contingency. But equally clear is that contingent changes at a genomic level often give rise to deterministic evolutionary responses at a phenotypic level, even at the level of transcriptomes that follows just above the genome in the hierarchical organization of organisms. Our results indicate that deterministic outcomes are not restricted to traits that evolve in convergence across many taxa and respond to prominent sources of selection. Instead, cryptic environmental variation—which is subsumed into categorical habitat types when we employ reductionist approaches—matters, because similar selective regimes beget similar evolutionary responses. Thus, the study of evolutionary repeatability and predictability has much to gain from multivariate approaches that reflect the complexity of organismal design and the multifarious nature of environmental gradients that exert selection on populations.

## Methods

The following sections provide a synopsis of the procedures used in this study. Detailed materials and methods are provided in the SI Appendix. Unless otherwise noted, analyses were conducted in R v. 4.1.1 (47).

### Sampling and RNA sequencing

Transcriptomes were reanalyzed from a previous study (17). In brief, gill samples preserved in RNAlater were collected from ten lineages that have independently colonized H_2_S-rich habitats in the United States, Mexico, and the Dominican Republic, as well as from ten geographically and phylogenetically proximate lineages that occupy nonsulfidic habitats (Table S1). Raw reads from transcriptome sequencing were mapped to the *Po. mexicana* reference genome [RefSeq accession number: GCF_001443325.1 (48)] with an appended mitochondrial genome (GenBank accession number: KC992998.1), which allowed for the generation of a read count matrix for each gene and individual.

### Phylogenelic analysis

The phylogenetic framework used in this study was established previously based on lineage-specific consensus sequences of 167 random genes (26), including sequences from *Fundulus heterclitus* [GenBank accession number: JXMV00000000.1 (49)] that was used as an outgroup. Genes were concatenated and partitioned according to the most likely model of DNA substitution and then used in maximum likelihood analyses using *RAxML* (50). We recovered a robust tree consistent with previous poeciliid phylogenies and bootstrap support that was generally > 99 %. The best-scoring maximum likelihood tree was used in subsequent analyses.

### Expression variance and evolution models

To identify convergent patterns in gene expression, we extracted the top 5000 expressed genes across all individuals and used them as dependent variables in expression variance and evolution (EVE) models (24, 25). These models identify genes with convergent expression shifts in a set of foreground lineages (sulfidic populations) relative to a set of background lineages (nonsulfidic populations). We subsampled the best-scoring maximum likelihood tree and the matrix of read counts such that independent EVE models could be run for each possible pair of sulfide spring lineages (total of 45 pairwise comparisons); *i.e*., each model included ten nonsulfidic lineages in the background and two sulfidic lineages in the foreground. EVE models implement an extended

Ornstein-Uhlenbeck process that incorporates within-species expression variance to test for branch-specific shifts in gene expression by comparing likelihoods that an expression parameter (θ_i_) for a given gene is shared between two groups of lineages versus θ_i_ for that gene being significantly different between the groups (25). For each gene, we employed a likelihood ratio test (LRT_θ_) contrasting the null hypothesis (θ_i_^sulfidic^ = θ_i_^nonsulfidic^) to the alternative hypothesis (θ_i_^sulfidic^ ≠ θ_i_^nonsulfidic^) using a *X*^2^ distribution to assess statistical significance. To account for multiple testing, we calculated FDR adjusted *P*-values using the Benjamini-Hochberg procedure (51). The number of genes with evidence for significant (FDR < 0.05) convergent expression shifts was then summed for each pair of sulfide spring lineages, providing a measure of the degree of convergence.

### Hypothesis testing

To test whether the degree of convergence in gene expression was associated with phylogenetic relatedness or ecological similarity between sulfide spring lineages, we first generated two predictor matrices. Phylogenetic relatedness was estimated by calculating phylogenetic distances between all lineage pairs based on the best-scoring maximum likelihood tree. To calculate ecological similarity, we compiled measurements of water quality (temperature, pH, specific conductively, dissolved oxygen, and H_2_S concentration) and downloaded 19 bioclimatic variables (28) based on the geographic location of each spring (Table S1). To visualize variation in environmental conditions among springs, we conducted a principal component analysis on all 24 variables using a correlation matrix. To obtain a metric of ecological dissimilarity for each pair of sulfide spring lineages, we *z*-transformed all variables and then calculated pairwise Euclidean distances in environment space (24 dimensions). These procedures yielded three matrices: a matrix describing the degree of convergence between all lineage pairs (number of genes with significant expression shifts), a matrix with the phylogenetic distance between all lineage pairs, and a matrix with the degree of ecological dissimilarity between all lineage pairs. We tested for a correlation between the degree of convergence and phylogenetic distance using a partial Mantel test conditioned on the ecological dissimilarity matrix. Similarly, we tested for a correlation between the degree of convergence and ecological dissimilarity using a partial Mantel test conditioned on the phylogenetic distance matrix. *P*-values were based on 100,000 permutations.

## Acknowledgements

We thank Lars Grønvold for help with statistical analyses, and John Coffin, Madison Nobrega, and Libby Wilson for feedback on the manuscript. This work was supported by grants from the NSF (IOS-1463720, IOS-1557795, IOS-1557860, and IOS-1931657) and the US Army Research Office (W911NF-15-1-0175, W911NF-16-1-0225) to JLK and MT.

## Data accessibility

Data and code associated with all analyses are available on GitHub (https://github.com/michitobler/degreesofconvergence). Sequence data are available at the National Center for Biotechnology Information (NCBI) under Bio-Project accession numbers PRJNA473350 and PRJNA608180.

## Notes

https://github.com/michitobler/degreesofconvergence

### Competing Interest Statement

The authors have declared no competing interest.

