## Supplementary material for "Ecology drives the degree of convergence in the gene expression of extremophile fishes": Online Supplementary Material

### Extended Methods

*Sampling*

Transcriptomes were reanalyzed from a previous study that conducted RNAseq on gill tissues (1). Gills were studied because they are in direct contact with environmental $H_2S$ (2), are the site of many physiological processes involved in the maintenance of homeostasis (3), and exhibit strong transcriptional responses to $H_2S$ exposure (4, 5). Gill tissues for transcriptome sequencing were collected from 10 lineages of poeciliid fishes that independently colonized sulfide springs in the United States, Mexico, and the Dominican Republic (Table S4). For comparison, we also included 10 closely related lineages from adjacent nonsulfidic habitats (Table S4). All fish were collected by using a seine, and the gills of adult females were extracted from both sides of the body immediately after capture. All tissues were preserved in RNAlater (Ambion Inc.) and stored at -20 °C upon return to the laboratory.

*Quantifying gene expression*

For transcriptome sequencing, we extracted total RNA extracted from pulverized tissue using the NucleoSpin RNA kit (Machery-Nagel, Düren, Germany). We then isolated mRNA with yhe NEBNext Poly(A) mRNA Magnetic Isolation Module (New England Biolabs, Inc., Ipswich, MA, USA) and constructed cDNA libraries with the NEBNext Ultra Directional RNA Library Prep Kit for Illumina (New England Biolabs, Inc., Ipswich, MA, USA). All cDNA libraries were individually barcoded, quantified using Qubit and an Agilent 2100 Bioanalyzer High Sensitivity DNA chip, and pooled in sets of 11-12 samples based on nanomolar concentrations. All libraries were sequenced on an Illumina HiSeq 2500 with paired-end 101 base pair (bp) reads at the Washington State University Spokane Genomics Core. All raw sequencing data are available at the National Center for Biotechnology Information (NCBI) under Bio-Project accession numbers PRJNA473350 and PRJNA608180.

Raw reads from Illumina sequencing were sorted by barcode, and we used *Trimgalore!* v. 0.4.0 (6) to remove adapters and low quality bases. Trimmed reads were then mapped to the *Po. mexicana* reference genome [RefSeq accession number: GCF_001443325.1 (7)] with an appended mitochondrial genome (GenBank accession number: KC992998.1) using *BWA-MEM* (8). Finally, we generated a matrix of read counts for each gene and individual using *cufflinks* v. 2.2.1 (9). We removed any genes that were not consistently expressed across all samples and then selected the 5,000 top genes across all lineages for subsequent analyses. These focal genes were annotated by comparing them to the entries in the human SWISSPROT database (critical E-value 0.001; access date 04/15/2017) using BLASTx (48) and retaining the top BLAST hit for each gene based on the top high-scoring segment pair.

*Phylogenetic analysis*

The phylogenetic framework used in this study was established previously based on lineage-specific consensus sequences of 167 random genes extracted from the RNAseq data (10), including sequences from *Fundulus heterclitus* [GenBank accession number: JXMV00000000.1 (11)] that were used as an outgroup. Genes were concatenated and partitioned according to the most likely model of DNA substitution, as determined by *jModelTest* 2 v. 2.1.10 (12), and then used in maximum likelihood analyses using *RAxML* v. 8.2.9 (13). *RAxML* analyses were conducted with the Rapid Bootstrap Algorithm with 1,000 bootstrap replicates, and we used a thorough maximum likelihood (ML) search with each partition assigned its own GTR + Γ + I model. We recovered a robust phylogenetic tree consistent with previous poeciliid phylogenies (14–16). Bootstrap support that was mostly >99 %. The only exceptions were the two populations of *G. sexradiata* and *G. eurystoma* (bootstrap support 60%), which represent a complex of recently diverged species (17).

*Expression variance and evolution models*

To identify convergent patterns in gene expression, we used the counts matrix of the top 5,000 genes and the best-scoring maximum likelihood tree as inputs in expression variance and evolution (EVE) models, as implemented in the *twoThetaTest()* function from the *evemodel* v. 0.0.0.9007 package in *R* v. 4.1.1 (18, 19). EVE models implement an extended Ornstein-Uhlenbeck process that incorporates within-species expression variance to test for branch-specific shifts in gene expression by comparing likelihoods that an expression parameter ($\theta_i$) for a given gene is shared between two groups of lineages versus $\theta_i$ for that gene being significantly different between the groups (19). We designated branches associated with lineages from sulfidic habitats as one group and those associated with nonsulfidic lineages as another group and then contrasted $\theta_i$ for each gene between these two groups. For each gene, we employed a likelihood ratio test ($LRT_\theta$) contrasting the null hypothesis ($\theta_i^{sulfidic} = \theta_i^{nonsulfidic}$) to the alternative hypothesis ($\theta_i^{sulfidic} \neq \theta_i^{nonsulfidic}$) using a $X^2$ distribution to assess statistical significance (19). To account for multiple testing, we calculated FDR adjusted *P*-values using the Benjamini-Hochberg procedure (20). EVE models were run in this manner for each possible pair of sulfide spring lineages (total of 45 pairwise comparisons). Hence, the counts matrix and the best-scoring maximum likelihood tree were subset, such that each model included ten nonsulfidic lineages in the background and two sulfidic lineages in the foreground. The number of genes with evidence for significant (FDR < 0.05) convergent expression shifts was then summed for each pair of sulfide spring lineages, providing a measure of the degree of convergence.

These models identify genes with convergent expression shifts in a set of foreground lineages (sulfidic populations) relative to a set of background lineages (nonsulfidic populations). We subsampled the best-scoring maximum likelihood tree and the matrix of read counts such that independent EVE models could be run; *i.e.,* EVE models implement an extended Ornstein-Uhlenbeck process that incorporates within-species expression variance to test for branch-specific shifts in gene expression by comparing likelihoods that an expression parameter ($\theta_i$) for a given gene is shared between two groups of lineages versus $\theta_i$ for that gene being significantly different between the groups. For each gene, we employed a likelihood ratio test ($LRT_\theta$) contrasting the null hypothesis ($\theta_i^{sulfidic} = \theta_i^{nonsulfidic}$) to the alternative hypothesis ($\theta_i^{sulfidic} \neq \theta_i^{nonsulfidic}$) using a $X^2$ distribution to assess statistical significance. To account for multiple testing, we calculated FDR adjusted *P*-values using the Benjamini-Hochberg procedure. The number of genes with evidence for significant (FDR < 0.05) convergent expression shifts was then summed for each pair of sulfide spring lineages, providing a measure of the degree of convergence.

*Phylogenetic and ecological distances*

To test whether the degree of convergence in gene expression was associated with phylogenetic relatedness or ecological similarity between sulfide spring lineages, we first generated two predictor matrices. Phylogenetic relatedness was estimated by calculating phylogenetic distances between all lineage pairs based on the best-scoring maximum likelihood tree using the *cophenetic*() function from the *stats* v. 3.6.2 package in R. To calculate ecological similarity, we assembled two sets of environmental data. First, we measured aspects of water quality in all sulfide springs. Temperature, pH, specific conductively, and dissolved oxygen were measured using a Hydrolab Multisonde 4A (Hach Environmental). Environmental $H_2S$ concentrations were measured using a methylene blue assay using a Hach DR1900 Portable Spectrophotometer (Hach Company, Loveland, CO, USA). Measurements and calibration of probes were conducted according to the manufacturer's recommendations. Measurements for each site represent averages from at least three Hydrolab readings and two $H_2S$ samples, although estimates for most springs spanned repeated visits over multiple years. Second, we also downloaded 19 bioclimatic variables (21) based on the geographic location of each spring. Specifically, we used the *getData*() function from the *raster* v. 3.5-2 package (22) to download bioclim variables from the WorldClim (21) repository at a spatial resolution of 0.5 minutes of a degree. Bioclim variables included annual mean temperature, mean diurnal range, isothermality, temperature seasonality, maximum temperature of warmest month, minimum temperature of coldest month, temperature annual range, mean temperature of wettest quarter, mean temperature of driest quarter, mean temperature of warmest quarter, mean temperature of coldest quarter, annual precipitation, precipitation of wettest month, precipitation of driest month, precipitation seasonality, precipitation of wettest quarter, precipitation of driest quarter, precipitation of warmest quarter, and precipitation of coldest quarter (see Table S3). To visualize variation in environmental conditions among springs, we conducted a principal component analysis with a correlation matrix on all 24 variables using the *prcomp*() function from the *stats* v. 3.6.2 package in R. To obtain a metric of ecological dissimilarity for each pair of sulfide spring lineages, we *z*-transformed all variables using the *scale*() function from base R and then calculated pairwise Euclidean distances in environmental space (24 dimensions) using the *dist*() function from the *stats* v. 3.6.2 package.

*Hypothesis testing*

To test hypotheses about the relationship between the degree of convergence, phylogenetic distance, and ecological distance, we used partial Mantel tests based on the matrices describing the degree of convergence between all lineage pairs (number of genes with significant expression shifts; square-root-transformed), phylogenetic distance between all lineage pairs, and the degree of ecological dissimilarity between all lineage pairs. We tested for a correlation between the degree of convergence and phylogenetic distance using a partial Mantel test conditioned on the ecological dissimilarity matrix. Similarly, we tested for a correlation between the degree of convergence and ecological dissimilarity using a partial Mantel test conditioned on the phylogenetic distance matrix. Partial Mantel tests were conducted using the *mantel*() function form the *ecodist* v. 2.0.7 package , and *P*-values were based on 100,000 permutations.

### Supplementary Figures

Figure S1. GO enrichment analysis of genes that exhibit a signature of convergent expression shifts in the majority ( ≥23) of lineage pairs. Note that this figure is provided as a separate file (FigureS1.png).

### Supplementary Tables

Table S1. List of sulfide springs included in this study, including their geographic locations, latitude and longitude, the fish lineages inhabiting them, water quality parameters measured in situ, and bioclimatic variables download from WorldClim.

| | **La Lluvia** | **El Azufre II** | **Baños del Azufre** | **La Gloria** | **Mogote del Puyacatengo** | **Green Springs** | **La Zurza** |
|---|---|---|---|---|---|---|---|
| Location | Chiapas, Mexico | Tabasco, Mexico | Tabasco, Mexico | Chiapas, Mexico | Tabasco, Mexico | Florida, USA | Dominican Republic |
| Lat/Long | 17.464/-92.895 | 17.438/-92.775 | 17.552/-92.999 | 17.532/-93.015 | 17.582/-92.900 | 28.863/-81.248 | 18.398/-71.570 |
| Lineage | *Po. mexicana* (Puy) | *Po. mexicana* (Tac) | *Po. mexicana* (Pic)<br>*G. eurystoma* | *Ps. bimaculatus*<br>*X. hellerii* | *G. sexradiata* | *Po. latipinna*<br>*G. holbrooki* | *L. sulphurophila* |
| DO [mg/l] | 1.73 | 1.29 | 1.06 | 0.99 | 1.31 | 0.6 | 4.8 |
| Temperature [°C] | 25.7 | 27.9 | 31.2 | 29.4 | 27.2 | 26.3 | 26.2 |
| pH | 7.2 | 6.8 | 6.8 | 6.8 | 6.9 | 7.1 | 7.3 |
| Specific conductivity [µScm$^{-1}$] | 2285 | 3992 | 2697 | 2238 | 1931 | 2645 | 1257 |
| $H_2S$ [µM/l] | 26.2 | 129.2 | 190.4 | 154.8 | 36.7 | 49.4 | 11.8 |
| Bio1: Annual mean temperature | 232 | 247 | 258 | 259 | 258 | 215 | 268 |
| Bio2: Mean diurnal range | 112 | 114 | 114 | 115 | 114 | 114 | 119 |
| Bio3: Isothermality | 66 | 66 | 65 | 64 | 65 | 46 | 77 |
| Bio4: Temperature seasonality | 1817 | 1851 | 1979 | 2030 | 1979 | 4733 | 1138 |
| Bio5: Max. temperature of warmest month | 316 | 335 | 348 | 351 | 348 | 328 | 342 |
| Bio6: Min. temperature of coldest month | 148 | 163 | 174 | 174 | 174 | 83 | 188 |
| Bio7: Temperature annual range | 168 | 172 | 174 | 177 | 174 | 245 | 154 |
| Bio8: Mean temperature of wettest quarter | 241 | 255 | 268 | 270 | 268 | 270 | 278 |
| Bio9: Mean temperature of driest quarter | 242 | 243 | 269 | 270 | 269 | 162 | 251 |
| Bio10: Mean temperature of warmest quarter | 250 | 266 | 278 | 280 | 278 | 270 | 281 |
| Bio11: Mean temperature of coldest quarter | 205 | 220 | 230 | 230 | 230 | 151 | 251 |
| Bio12: Annual precipitation | 3579 | 3189 | 3478 | 3637 | 3478 | 1306 | 842 |
| Bio13: Precipitation of wettest month | 518 | 493 | 535 | 553 | 535 | 184 | 136 |

| | | | | | | | |
|---|---|---|---|---|---|---|---|
| Bio14: Precipitation of driest month | 125 | 112 | 114 | 116 | 114 | 54 | 23 |
| Bio15: Precipitation seasonality | 43 | 45 | 45 | 46 | 45 | 45 | 54 |
| Bio16: Precipitation of wettest quarter | 1407 | 1292 | 1411 | 1491 | 1411 | 538 | 335 |
| Bio17 Precipitation of driest quarter | 439 | 397 | 386 | 386 | 386 | 176 | 72 |
| Bio18 Precipitation of warmest quarter | 689 | 641 | 800 | 847 | 800 | 531 | 266 |
| Bio19 Precipitation of coldest quarter | 687 | 583 | 701 | 739 | 701 | 206 | 72 |

Table S2. Summary of EVE results. The table includes a list of all 5,000 genes included in the analyses, their gene ID, SwissProt accession number and annotation, the number of models in which the gene exhibited a convergent expression shift, and the FDR-corrected *P*-value for each model. Note that this table is included as a separate spreadsheet (TableS2.csv).

Table S3. Results of the principal component analysis of environmental variables.

| **Variable** | **PC1** | **PC2** | **PC3** |
|---|---|---|---|
| Dissolved oxygen | 0.057 | 0.322 | -0.223 |
| **Temperature** | **-0.217** | 0.004 | 0.299 |
| **pH** | **0.227** | 0.097 | -0.276 |
| Specific conductivity | -0.045 | -0.235 | 0.053 |
| **$H_2S$ concentration** | **-0.208** | -0.053 | 0.287 |
| **Bio1: Annual mean temperature** | **-0.203** | 0.259 | 0.034 |
| Bio2: Mean diurnal range | 0.066 | 0.338 | 0.117 |
| Bio3: Isothermality | -0.161 | 0.260 | -0.224 |
| **Bio4: Temperature seasonality** | **0.204** | -0.211 | 0.220 |
| Bio5: Max. temperature of warmest month | -0.178 | 0.170 | 0.336 |
| **Bio6: Min. temperature of coldest month** | **-0.225** | 0.223 | -0.085 |
| **Bio7: Temperature annual range** | **0.204** | -0.206 | 0.231 |
| Bio8: Mean temperature of wettest quarter | 0.063 | 0.202 | 0.429 |
| **Bio9: Mean temperature of driest quarter** | **-0.261** | 0.154 | -0.064 |
| Bio10: Mean temperature of warmest quarter | -0.063 | 0.212 | 0.417 |
| **Bio11: Mean temperature of coldest quarter** | **-0.209** | 0.246 | -0.079 |
| **Bio12: Annual precipitation** | **-0.273** | -0.111 | -0.067 |
| **Bio13: Precipitation of wettest month** | **-0.279** | -0.095 | -0.051 |
| **Bio14: Precipitation of driest month** | **-0.256** | -0.162 | -0.087 |
| Bio15: Precipitation seasonality | 0.084 | 0.344 | 0.027 |
| **Bio16: Precipitation of wettest quarter** | **-0.274** | -0.110 | -0.048 |
| **Bio17 Precipitation of driest quarter** | **-0.252** | -0.164 | -0.113 |
| **Bio18 Precipitation of warmest quarter** | **-0.241** | -0.164 | 0.116 |
| **Bio19 Precipitation of coldest quarter** | **-0.273** | -0.113 | -0.035 |
| Eigenvalue | 3.432 | 2.751 | 1.833 |
| Proportion of variance | 0.491 | 0.315 | 0.140 |

Table S4. Locality information of all samples included in this study.

| **Species** | **Locality** | **Lat/Long** |
|---|---|---|
| *Sulfidic lineages* | | |
| *Poecilia mexicana* (Puy) | La Lluvia springs, Rio Puyacatengo drainage, Tabasco, MX | 17.464/-92.895 |
| *Poecilia mexicana* (Tac) | El Azufre, Rio Tacotalpa drainage, Tabasco, MX | 17.438/-92.775 |
| *Poecilia mexicana* (Pic) | Baños del Azufre, Rio Pichucalco drainage, Tabasco, MX | 17.552/-92.999 |
| *Poecilia latipinna* | Green Springs, St. Johns River drainage, Florida, USA | 28.863/-81.248 |
| *Limia sulphurophila* | Balnearios La Zurza, Lago Enriquillo basin, Independencia, DR | 18.398/-71.570 |
| *Gambusia sexradiata* | Mogote del Puyacatengo, Rio Puyacatengo drainage, Tabasco, MX | 17.582/-92.900 |
| *Gambusia eurystoma* | Baños del Azufre, Rio Pichucalco drainage, Tabasco, MX | 17.552/-92.999 |
| *Gambusia holbrooki* | Green Springs, St. Johns River drainage, Florida, USA | 28.863/-81.248 |
| *Pseudoxiphophorus bimaculatus* | La Gloria springs, Rio Pichucalco drainage, Chiapas, MX | 17.532/-93.015 |
| *Xiphophorus hellerii* | La Gloria springs, Rio Pichucalco drainage, Chiapas, MX | 17.532/-93.015 |
| *Nonsulfidic lineages* | | |
| *Poecilia mexicana* | Arroyo Bonita, Rio Tacotalpa drainage, Tabasco, MX | 17.427/-92.752 |
| *Poecilia mexicana* | Arroyo Rosita, Rio Pichucalco drainage, Chiapas, MX | 17.485/-93.104 |
| *Poecilia mexicana* | Rio Puyacatengo, Rio Puyacatengo drainage, Tabasco, MX | 17.510/-92.914 |
| *Poecilia limantouri* | Rio Garces, Rio Panuco drainage, Hidalgo, MX | 20.940/-98.282 |
| *Poecilia latipinna* | Mariner's Cove, Lake Monroe, St. Johns River drainage, Florida, USA | 28.857/-81.239 |
| *Limia perugiae* | Stream in Cabral, Rio Yaque del Sur drainage, Barahona, DR | 18.246/-71.223 |
| *Gambusia sexradiata* | Laguna Sitio Grande, Rio Ixtapangajoya drainage, Tabasco, MX | 17.677/-92.997 |
| *Gambusia holbrooki* | Mariner's Cove, Lake Monroe, St. Johns River drainage, Florida, USA | 28.857/-81.239 |
| *Pseudoxiphophorus bimaculatus* | Arroyo Pujil, Rio Ixtapangajoya drainage, Chiapas, MX | 17.476/-92.986 |
| *Xiphophorus hellerii* | Rio El Azufre, west branch, Rio Pichucalco drainage, Chiapas, MX | 17.556/-93.008 |
