## Supplementary figures and images for "Ecology drives the degree of convergence in the gene expression of extremophile fishes"

### Figure S1

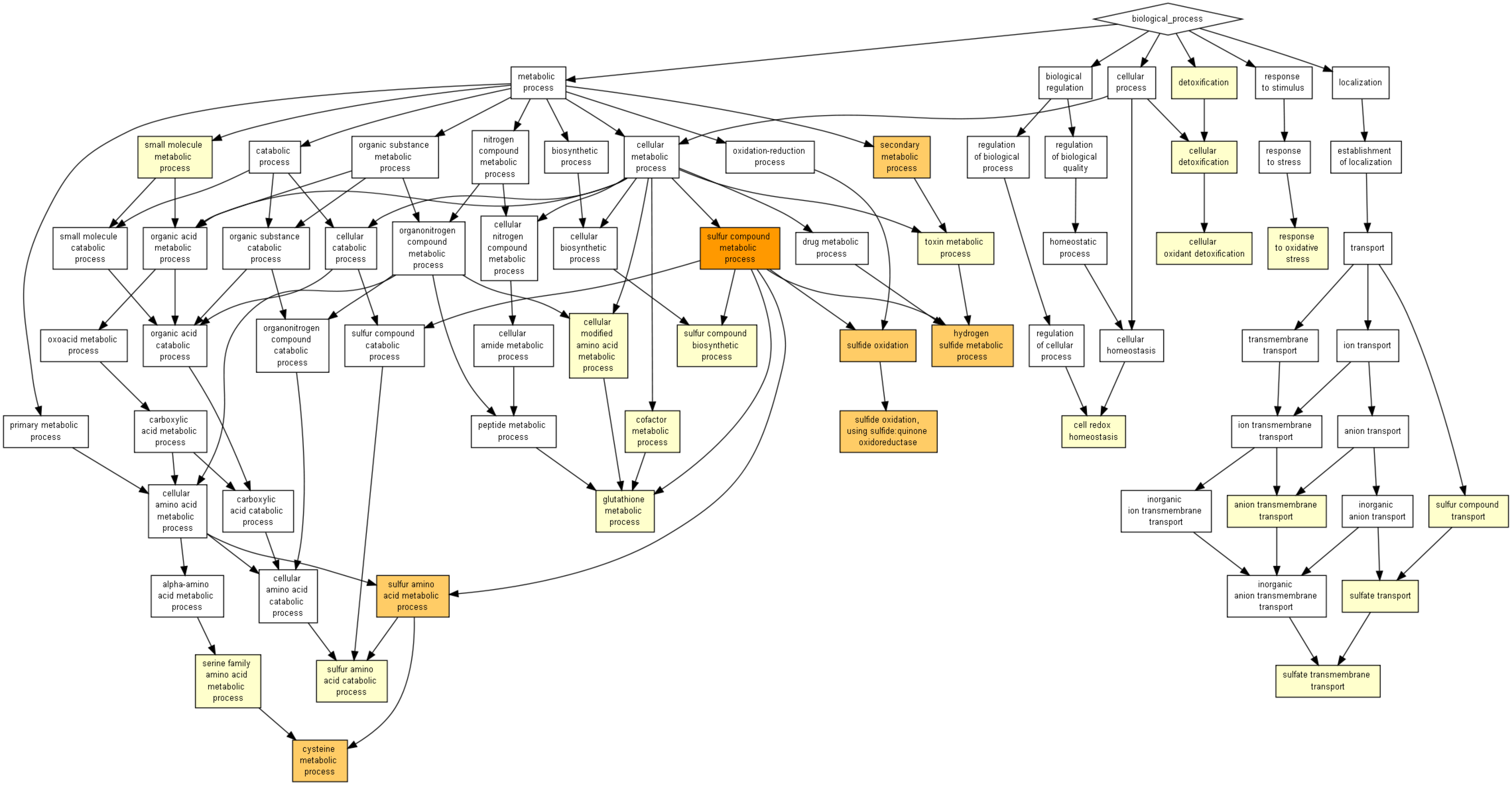
